# Phosphatidylinositol 4,5 bisphosphate controls the cis and trans interactions of synaptotagmin 1

**DOI:** 10.1101/582965

**Authors:** S.B. Nyenhuis, A. Thapa, D. S. Cafiso

## Abstract

Synaptotagmin 1 acts as the Ca^2+^-sensor for synchronous neurotransmitter release; however, the mechanism by which it functions is not understood and is presently a topic of considerable interest. Here we describe measurements on full-length membrane reconstituted synaptotagmin 1 using site-directed spin labeling where we characterize the linker region as well as the *cis* (vesicle membrane) and *trans* (cytoplasmic membrane) binding of its two C2 domains. In the full-length protein, the C2A domain does not undergo membrane insertion in the absence of Ca^2+^; however, the C2B domain will bind to and penetrate in *trans* to a membrane containing phosphatidylinositol 4,5 bisphosphate (PIP_2_), even if phosphatidylserine (PS) is present in the cis membrane. In the presence of Ca^2+^, the Ca^2+^-binding loops of C2A and C2B both insert into the membrane interface; moreover, C2A preferentially inserts into PS containing bilayers and will bind in a *cis* configuration to membranes containing PS even if a PIP_2_ membrane is presented in *trans*. The data are consistent with a bridging activity for Syt1 where the two domains bind to opposing vesicle and plasma membranes. The failure of C2A to bind membranes in the absence of Ca^2+^ and the long unstructured segment linking C2A to the vesicle membrane indicates that synaptotagmin 1 could act to significantly shorten the vesicle-plasma membrane distance with increasing levels of Ca^2+^.

---

Neuronal exocytosis involves a highly coordinated series of molecular steps that result in the fusion of the synaptic vesicle with the presynaptic membrane and the release of neurotransmitter into the synapase. The assembly of three soluble NSF attachment receptor proteins (SNAREs) into a four-helical bundle provides the energy to drive this fusion event (1, 2), but the sequence of assembly of these SNAREs and the mechanism by which increased Ca^2+^ levels trigger SNARE assembly are not known. The three neuronal SNAREs include syntaxin 1A and SNAP25 in the plasma membrane, and synaptobrevin in the vesicle membrane (3-5), and they appear to interact with and be regulated by Munc18, Munc13 and complexin (6-9). The binding of Ca^2+^ to Synaptotagmin 1 (Syt1) triggers synchronous neurotransmitter release and is the primary Ca^2+^ sensing protein in this system (10).

Synaptotagmin 1 is anchored to the synaptic vesicle membrane by a single transmembrane helix. As shown in Fig. 1, a long linker of about 60 residues connects the transmembrane helix to two Ca^2+^-binding C2 domains that are connected by a short flexible 6 residue segment (11). The binding of Ca^2+^ to Syt1 increases the affinity of the C2 domains towards membranes containing negatively charged lipids and results in the interfacial insertion of the Ca^2+^ binding loops of the domains. In the absence of Ca^2+^, the C2A domain has no significant membrane affinity, although C2B has a weak affinity to membranes containing phosphatidylserine (PS), and a stronger affinity to membranes with phosphatidylinositol 4,5 bisphosphate (PIP_2_) (12). This Ca^2+^-independent binding mode of the C2B domain is mediated by its highly positively charged polybasic face (13).

**Figure 1.**
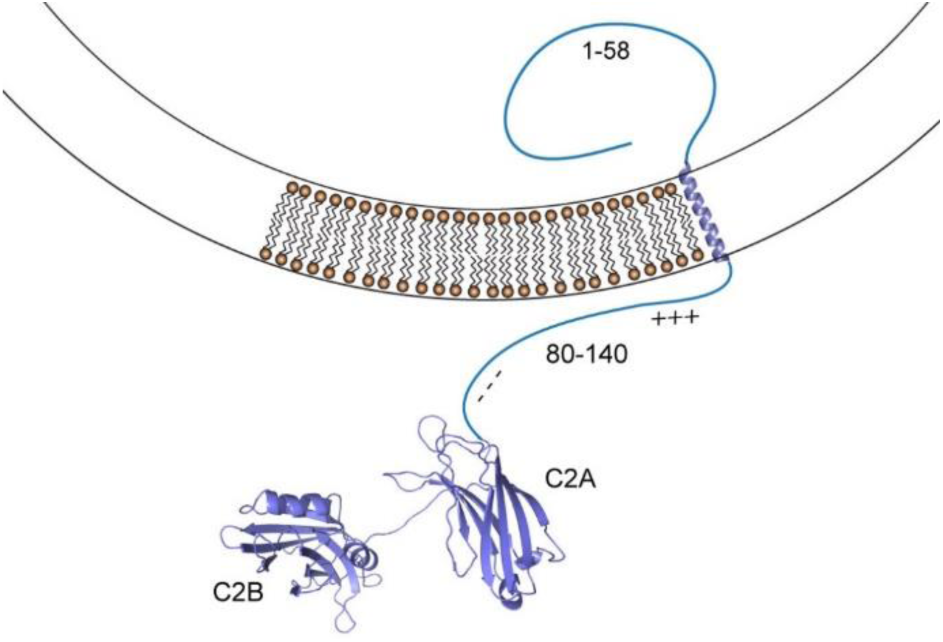
Syt1 is a synaptic vesicle associated membrane proteins with two C2 domains. The domains are linked by a long, 60 residue segment that connects C2A to a transmembrane helix. Regions of the linker near the membrane are positively charged, while segments closer to C2A are negatively charged.

In addition to binding membranes, Syt1 also shows an affinity towards SNAREs. Several different structural models for Syt1-SNARE interactions have been generated using NMR and modeling (14), crystallography (15, 16) and smFRET (17). These models indicate that multiple Syt1-SNARE interactions are possible, but because they involve interactions with assembled SNARE complexes that represent a post-fusion state (18), they do not reveal how Syt1 might regulate fusion. Moreover, Syt1-SNARE interactions appear to be weak and heterogeneous and are diminished or eliminated when ionic compositions resembling those in the cell are present (19). Under these same conditions, Ca^2+^-dependent interactions of Syt1 to membranes, particularly to membranes containing PIP_2_ or a combination of PS and PIP_2_, are maintained.

These observations suggest that while Syt1-SNARE interactions might help localize Syt1 near the SNAREs, they are unlikely to be directly responsible for Ca^2+^-mediated synchronous release. The most robust interactions appear to be to membranes, and a number of models have been proposed that involve the regulation of fusion through the membrane interactions of Syt1. For example, Syt1 has been found to demix PS from the bulk lipid and to alter the acyl chain order of the bilayer in a Ca^2+^-dependent manner (20). Acyl chain order has been shown to alter the structure of the SNAREs and Syt1 has been shown to alter the orientation of the SNAREs in a Ca^2+^-dependent manner (21). These observations suggest a model where Syt1 triggers fusion by acting through the membrane and placing the SNAREs in a fusion competent state. Synaptotagmin 1 has also been shown to alter the curvature properties of bilayers; and in a second model, Syt1 is proposed to modify curvature in a Ca^2+^-dependent manner and act with the SNAREs to overcome the energetic barriers to fusion (22). Earlier work demonstrated that the two C2 domains of Syt1 have the capacity to bridge between two bilayers (23, 24), and there is evidence that Syt1 may act to significantly shorten the vesicle-plasma membrane distance in the presence of Ca^2+^ (25). This suggests a model where changes in vesicle-plasma membrane distance might place the pre-fusion SNARE complex or its regulatory partners in an active configuration. Finally, a quite different model has been proposed that involves the oligomerization of the C2B domains of Syt1 to form a large ring. This ring acts to sterically block interactions between the synaptic vesicle and the plasma membranes and the binding of Ca^2+^ to Syt1 results in the membrane insertion of the domains and the release of this steric inhibition (26).

The bulk of the structural work on Syt1 has been carried out on a soluble fragment of Syt1 containing its two C2 domains (sC2AB). In the present work, we examine the configuration and membrane insertion of the C2A and C2B domains in the context of full-length Syt1 (fSyt1). We identify conditions that lead to a bridging between two bilayers by fSyt1 and we characterize interactions within the long juxta-membrane linker of Syt1. We find no evidence that Syt1 oligomerizes in its full-length membrane associated form; however, Syt1 is ideally configured to regulate the vesicle-plasma membrane distance. In the absence of Ca^2+^, the C2B domain is likely to target the PIP_2_ containing plasma membrane while the C2A domain is free in the synaptic space along with 50 residues of the long juxta-membrane linker. In the presence of Ca^2+^, both domains insert into the bilayer. However, because their lipid preferences are different, the two domains are likely to bridge between the vesicle and plasma membranes, thereby shortening this interbilayer space.

## Materials and Methods

### Expression and Purification of Synaptotagmin 1 constructs

The C2B plasmid was provided by Jose Rizo-Rey (Texas Southwestern Medical Center) and expression and purification of sC2AB (residues 136-421), C2B (residues 271–421), and fSyt1 (residues 1–421) from *R. norvegicus* was carried out as previously published (12, 27, 28). The native cysteine at residue 277 was mutated to an alanine and sC2AB and sC2B were expressed using pGEX-KG constructs with N-terminal GST tags. For sedimentation assays, site E269 was mutated to a cysteine using PIPE site-directed mutagenesis to attach a fluorophore. The native cysteines (C73A, C74A, C76A, C78A, C82S, and C277S) were mutated in fSyt1 using the PIPE site-directed mutagenesis (28) and expressed in a pET-28a vector with an N-terminal HIS6-tag. For EPR measurements, single cysteine mutations were introduced into the cysteine-free construct at positions: K86C, K90C, K95C, M173C, V304C. Double mutants were introduced at sites K86C/P136C, P136C/L323C, M173C/V304C, M173C/L323C, M173C/A395C. DNA sequencing for all mutations were verified by Genewiz DNA sequencing (South Plainfield, NJ).

The sC2AB and C2B constructs were expressed in BL21(DE3) cells (Invitrogen, Carlsbad, CA) and then purified using GST affinity chromatography. For NMR experiments, C2B was grown in minimal media with ^15^NH_4_Cl (Cambridge Isotopes, Andover, MA) as the sole nitrogen source. GST tags were removed via thrombin cleavage, and the protein constructs were further purified by ion-exchange chromatography to remove any additional protein or nucleic acid contamination. The C2B domain was further purified with a final round of gel filtration chromatography. For binding measurements, sC2AB was fluorophore labelled at room temperature by first incubating with a 10-fold excess of DTT for 2 hours, desalting, labeling with Bodipy FL C_1_-IA for 2 hours, and finally running the protein through a PD10 column. Labeled protein was dialyzed against 12 L of 20 mM MOPS, 150 mM KCl, pH 7 to remove free label.

The fSyt1 construct was expressed in BL21(DE3)-RIL cells (Invitrogen, Carlsbad, CA) and purified in CHAPS using nickel affinity chromatography following a protocol described earlier (29). The fSyt1 was spin labeled with the thiol-specific spin label, MTSSL ((1-oxy-2,2,5,5-tetramethylpyrrolinyl-3-methyl)methanethiosulfonate), while bound to the nickel column overnight at 4°C. The HIS tag was removed via thrombin cleavage and fSyt1 was further purified by ion-exchange chromatography to remove any additional protein or nucleic acid contaminants. Full-length Syt1 was reconstituted into either: palmitoyloleoylphosphatidyl-choline (POPC):cholesterol (Chol) (80:20) or POPC:palmitoyloleoylphosphatidylserine (POPS (85:15) at a 1:200 protein to lipid ratio by dialysis in the presence of Bio-Beads (Bio-Rad, Hercules, CA) into metal free buffer. The purity of all constructs was verified by SDS–PAGE and the UV absorbance ratio (260/280 nm). The protein concentration was determined from the absorbance at 280 nm. All lipids were obtained from Avanti Polar Lipids (Alabaster, AL).

### Vesicle sedimentation binding assays

Sedimentation binding assays were performed as described previously (12, 30). In one assay, sucrose loaded 100 nm large unilamellar vesicles (LUVs) were titrated into Biodipy labeled sC2AB at a 2 nM concentration, and in a second assay, the lipid concentration of LUVs was maintained at 1 mM and ATP/Mg^2+^ or P_2_O_7_^4-^/Mg^2^ were increased in concentration to determine their effect upon sC2AB binding. Protein and LUVs were incubated for 10 minutes and the LUVs were pelleted at 100,000 x g, 20°C for 1 hour. Unbound protein in the supernatant was measured from Bodipy fluorescence emission (480 nm excitation and 520 nm emission) using a FluorMax3 (Jobin Yvon, Edison, NJ). Values were used to calculate the fraction of membrane-bound protein (*f*_*b*_) and the data fit to the expression *f*_*b*_ = *K*[*L*]/(1 + *K*[*L*]), where K is the reciprocal molar partition coefficient and [L] is the accessible lipid concentration. A standard phosphate assay was used to determine lipid concentrations (31, 32).

### Proton-nitrogen HSQC NMR

NMR experiments were performed on a Bruker 600 MHz NMR spectrometer as described previously (12), but using a shorter construct containing residues 271-421 (C2B). The NMR spectrum from 400 μM C2B was titrated against IP_3_ or ATP/Mg^2+^. The protein was in NMR buffer (50 mM MES, 150 mM NaCl, 3 mM CaCl_2_, 10% D_2_O, pH 6.3) in Shigemi tubes and the spectrum acquired at 27°C for 2 hours. New protein samples were used for each titration point to avoid precipitation of C2B. The NMR data were processed in NMRPipe (33), and the residue assignments were matched in Sparky from the PDB NMR resonance assignments of C2B (PDB 1K5W) (27, 34). Chemical shift analysis was performed using CCPN NMR Analysis V2 (35). To determine the apparent affinity of ATP and IP_3_ to C2B, chemical shifts for each residue were fit using a standard binding isotherm: *y* = *a*(*bx*/(1 + *bx*). Images for the chemical shift and binding constants were plotted onto the C2B domain using Pymol (36).

### Continuous Wave EPR

Experiments were performed as stated previously on a CW X-Band EMX spectrometer (Bruker Biospin, Billerica, MA) equipped with an ER 4123D dielectric resonator (28). All EPR spectra were recorded at room temperature with 100-G magnetic field sweep, 1-G modulation, and 2.0-milliwatt (mW) incident microwave power. The measurements were performed on 6 μl samples in glass capillary tubes (0.60 mm inner diameter × 0.84 mm outer diameter round capillary; VitroCom, Mountain Lakes, NJ). The phasing, normalization, and subtraction of EPR spectra were performed using LabVIEW software provided by Dr. Christian Altenbach (UCLA, Los Angeles, CA) or with in lab software written by David Nyenhuis.

To assess the *cis* or *trans* binding of fSyt1, EPR spectra from R1 labeled fSyt1 were recorded in reconstituted proteoliposomes using either negatively charged (POPC:POPS (85:15)) or uncharged lipid compositions (POPC:Chol (80:20)). The protein was reconstituted at a protein:lipid ratio of 1:200 and measured at a concentration of 50 μM protein. To determine *trans* binding, LUVs containing charged lipids at a 10 mM concentration (POPC:POPS (90:10) or POPC:PIP_2_ (95:5)) were added to the reconstituted fSyt1. The EPR spectra were recorded in a metal free state, in the presence of 1 mM Ca^2+^, after the addition ATP/Mg^2+^, and then following the addition of 4 mM EGTA. Progressive Power saturation of the EPR spectra was performed as described previously (28, 37). Samples were placed in gas permeable TPX-2 capillaries, and each sample run in the presence of air (O_2_) or Ni(II)EDDA to determine the membrane depth parameter, Φ. The spin label depth was estimated using the expression: Φ = *A*[*tanh*(*B*(*x* − *C*))] + *D* where, *x* is the distance of the spin label from the phospholipid phosphate plane in the bilayer and *A, B, C* and *D* are empirically determined constants (28, 37).

### Pulsed EPR

Experiments were performed on a Bruker Elexsys E580 EPR spectrometer running at Q-band using an EN5107D2 dielectric resonator. Measurements were performed on single or double labeled fSyt in 20% deuterated glycerol at a concentration of about 100-200 μM using a 12 μL sample volume. Samples were run in physiological buffer (25 mM HEPES, 150 mM KCl, pH 7.4) with or without the addition of calcium, unless otherwise stated, high salt buffer (25 mM HEPES, 300 mM KCl, pH 7.4) or low salt buffer (25 mM HEPES, 25 mM KCl, pH 7.4). Samples were loaded into quartz capillaries of 1.5 ID x 1.8 OD 100 mm length and flash frozen in liquid nitrogen then run at 80 K. DEER data were collected using a dead-time free four-pulse sequence with 16-step phase cycling (one 16 ns π/2 and two 32 ns π observe pulses separated by a 32 ns π pump pulse).(38, 39) Pump and observe pulses were separated by 75 MHz. DeerAnalysis2015 or DD was used for the removal of the dipolar background from the raw DEER data, V(t)/V(0) (39-42), and Tikhonov regularization was used to extract distance distributions from the resulting form factors, F(t)/F(0) (38, 43).

## Results

### Polyelectrolytes at low mM concentrations interfere with the Ca^2+^-dependent binding of Syt1 C2AB to bilayers containing PS but not to membranes containing PIP_***2***_

Moderate levels of ATP interfere with the Ca^2+^-dependent binding of reconstituted fSyt1 to vesicles containing phoshatidylserine (PS), but not to vesicles containing PIP_2_ (44). To explore the effect of ATP on the membrane binding affinity of Syt1, we used the soluble fragment of Syt1 containing its two C2 domains (sC2AB), to quantitate the Ca^2+^-dependent binding of sC2AB to membranes using a vesicle sedimentation assay (30). The equilibrium binding of sC2AB in the presence of Ca^2+^ to membranes containing 15 mol% POPS is shown in Fig. 2a, and yields a partition coefficient of approximately 2.2±0.4 x 10^4^ M^-1^, indicating that 50% of the protein is membrane associated at an accessible lipid concentration of about 50 μM. This is within the values obtained in previous sedimentation measurements (12). As seen in Fig. 2a, the addition of 1mM pyrophosphate (P_2_O_7_^4-^/Mg^2+^) reduces the affinity of sC2AB approximately 3-fold, and the addition of 1 mM ATP/Mg^2+^ eliminates the membrane binding of sC2AB.

**Figure 2.**
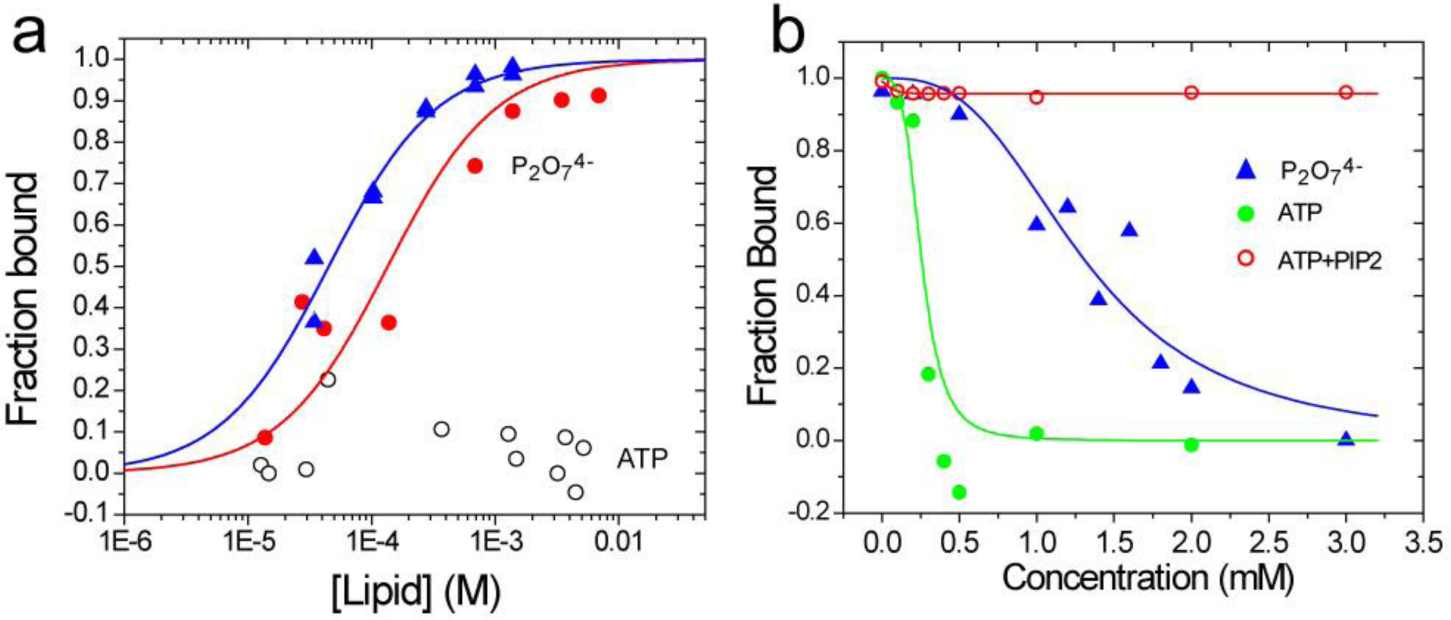
(**a**) Vesicle sedimentation assays of sC2AB in the presence of 1 mM Ca^2+^ yield the fraction sC2AB bound to POPC:POPS (85:15) LUVs as a function of the accessible lipid concentration (blue), upon the addition of 1 mM pyrophosphate/2 mM Mg^2+^ (red) and with the addition of 1 mM ATP/2 mM Mg^2+^ (open circles). (**b**) At a POPC:POPS (85:15) lipid concentration of 1 mM, sC2AB is almost fully membrane associated in the presence of Ca^2+^. The fraction of membrane associated sC2AB at this lipid concentration is shown as a function of added ATP/Mg^2+^ (green) or pyrophosphate/Mg^2+^ (blue). The fraction of bound sC2AB to 1 mM POPC:POPS:PIP_2_ (84:15:1) with added ATP/Mg^2+^ is shown in the red trace.

In Fig. 2b, the effect of adding either ATP/Mg^2+^ or P_2_O_7_^4-^/Mg^2+^ on the partitioning of sC2AB to POPC:POPS (85:15) LUVs was examined. We fixed the lipid concentration at 1 mM, which is sufficient to bind virtually all sC2AB (Fig. 2a). As seen in Fig. 2b, the membrane binding of sC2AB is eliminated by a few hundred μM ATP/Mg^2+^ or 2 to 3 mM P_2_O_7_^4-^/Mg^2+^. However, when 1 mol% PIP_2_ is added to the POPC:POPS mixture (red trace), ATP no longer displaces sC2AB from the membrane interface even up to levels of 3 mM.

In the full-length protein, the membrane insertion of the C2 domains was examined by incorporating the spin labeled R1 side chain into the 1^st^ Ca^2+^-binding loop of either the C2A or C2B domains of membrane reconstituted fSyt1. These loops are known to undergo Ca^2+^-dependent membrane insertion in sC2AB (45) as well as fSyt1 (28, 29) and Fig. 3a shows the EPR spectra obtained from sites 173R1 and 304R1 when fSyt1 is reconstituted into membranes containing POPC:POPS (85:15). The dramatic broadening in the EPR lineshape upon Ca^2+^ addition is due to insertion into the membrane environment, which limits the rate of interconversion between R1 rotamers (46, 47). Membrane insertion can also be confirmed by progressive power saturation of the EPR spectrum (48), where membrane depths of the R1 side chain attached to sC2AB and fSyt1 have been obtained (see Table S1) (12, 28).

**Figure 3.**
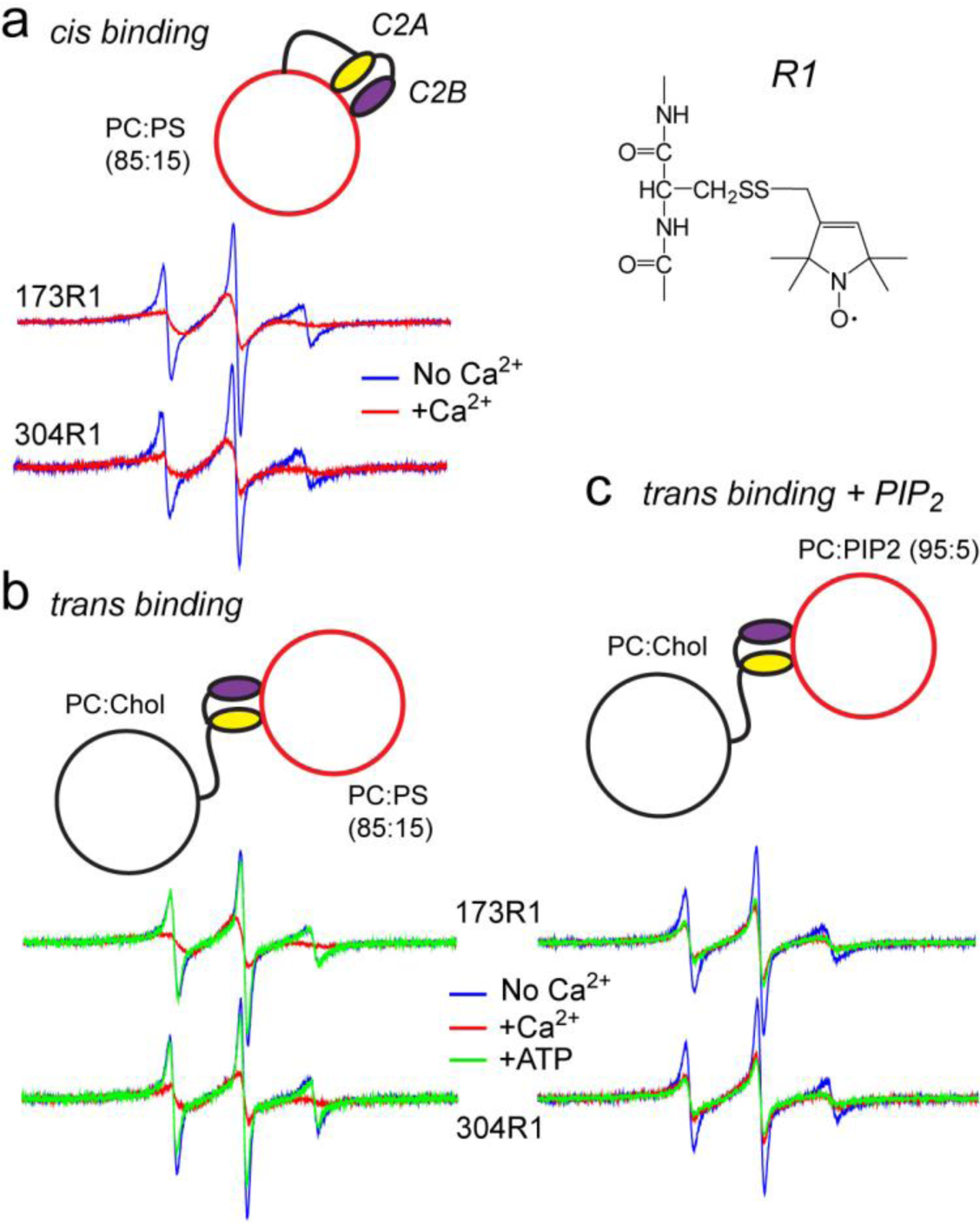
EPR spectra from C2A and C2B reflect *cis* or *trans* membrane insertion of these domains. **(a)** EPR spectra from sites 173 and 304 in fSyt1 labeled with the spin labeled side chain, R1, when reconstituted into PC:PS vesicles in the absence (blue trace) and presence (red trace) of 1 mM Ca^2+^. **(b)** EPR spectra from fSyt1 when reconstituted into PC:Chol (80:20) in the presence of vesicles of PC:PS (85:15) at a lipid concentration of 10 mM. Both domains insert in the presence of Ca^2+^ (red trace), but the addition of 1 mM ATP/Mg^2+^ eliminates the membrane insertion (green trace). (**c**) EPR spectra from fSyt1, when lipid vesicles composed of PC:PIP_2_ (95:5) are added. The addition of Ca^2+^induces membrane penetration of either domain, but 1 mM ATP/Mg^2+^ does not reverse the interaction.

The case shown in Fig. 3a is likely to result from the binding of C2A and C2B to the same (*cis*) vesicle into which Syt1 is reconstituted. However, binding in *trans* to a second membrane vesicle can be observed if negatively charged lipid is eliminated from the *cis* vesicle. When incorporated into uncharged vesicles, the EPR spectra show no evidence for a *cis* membrane insertion of C2A or C2B with or without Ca^2+^, and this lack of insertion is also evident from power saturation of the EPR spectrum (see Katti et al.(28) and Table S1). However, when PC:PS (85:15) vesicles are added to this suspension of neutral membranes containing labeled fSyt1 (Fig. 3b), a Ca^2+^-dependent *trans* insertion is observed for both the C2A and C2B domains. The EPR spectra with Ca^2+^ in Fig. 3b are virtually identical to those seen in Fig. 3a. In this case, addition of 1 mM ATP/Mg^2+^ reverses the *trans* insertion of fSyt1. When the target membrane contains PC:PIP_2_ (95:5) (Fig. 3c), *trans* insertion also takes place upon Ca^2+^ addition, but insertion of the domains is not reversed by ATP/Mg^2+^. Unlike the case seen in Fig. 3b, ATP cannot reverse the POPC:POPS membrane insertion seen in Fig. 3a for C2A or C2B associating in a *cis* configuration. The inability to reverse this *cis* binding is likely due to a strong membrane affinity due to the locally high-concentration of negatively charged lipid experienced by the C2 domains on their proximal surface. This suggests that the Ca^2+^-dependent membrane binding seen in Fig. 3a for C2A and C2B is likely to be dominated by a *cis* interaction.

### Polyelectrolytes associate strongly with the polybasic face of C2B

Why does ATP interfere with sC2AB membrane binding? In these experiments, the levels of ATP added are not sufficient to chelate the available Ca^2+^ (49). However, acidic contaminants such as nucleic acids are known to strongly associate with the C2B domain on its highly polybasic surface, and this contamination is well known to modify the behavior of C2B (50) (24). The inhibition of the Syt1 membrane interaction by ATP/Mg^2+^ and the ability of PIP_2_ to overcome this inhibition is likely due to a competition between ATP/Mg^2+^ and PIP_2_ for the polybasic face of C2B. Shown in Fig. 4 are chemical shift maps obtained from HSQC spectra of C2B in the presence of either IP_3_ (the headgroup of PIP_2_) or ATP (Figures S1-S3). The chemical shifts produced in C2B by either of these polyanions are localized in and near the polybasic face. Shown in Fig. 4b are the relative affinities determined from the chemical shifts when either ATP or IP_3_ are titrated into C2B. The average affinities of IP_3_ were found to be approximately 16 μM, while the affinities of ATP were about 20-fold weaker, or approximately 320 μM. The construct used in these experiments is important, as the affinity of IP_3_ measured here is substantially stronger than that reported earlier for a longer C2B fragment (12). The relative differences in affinity between IP_3_ and ATP are consistent with the observation that PIP_2_ can overcome the inhibition in Syt1 membrane binding produced by ATP.

**Figure 4.**
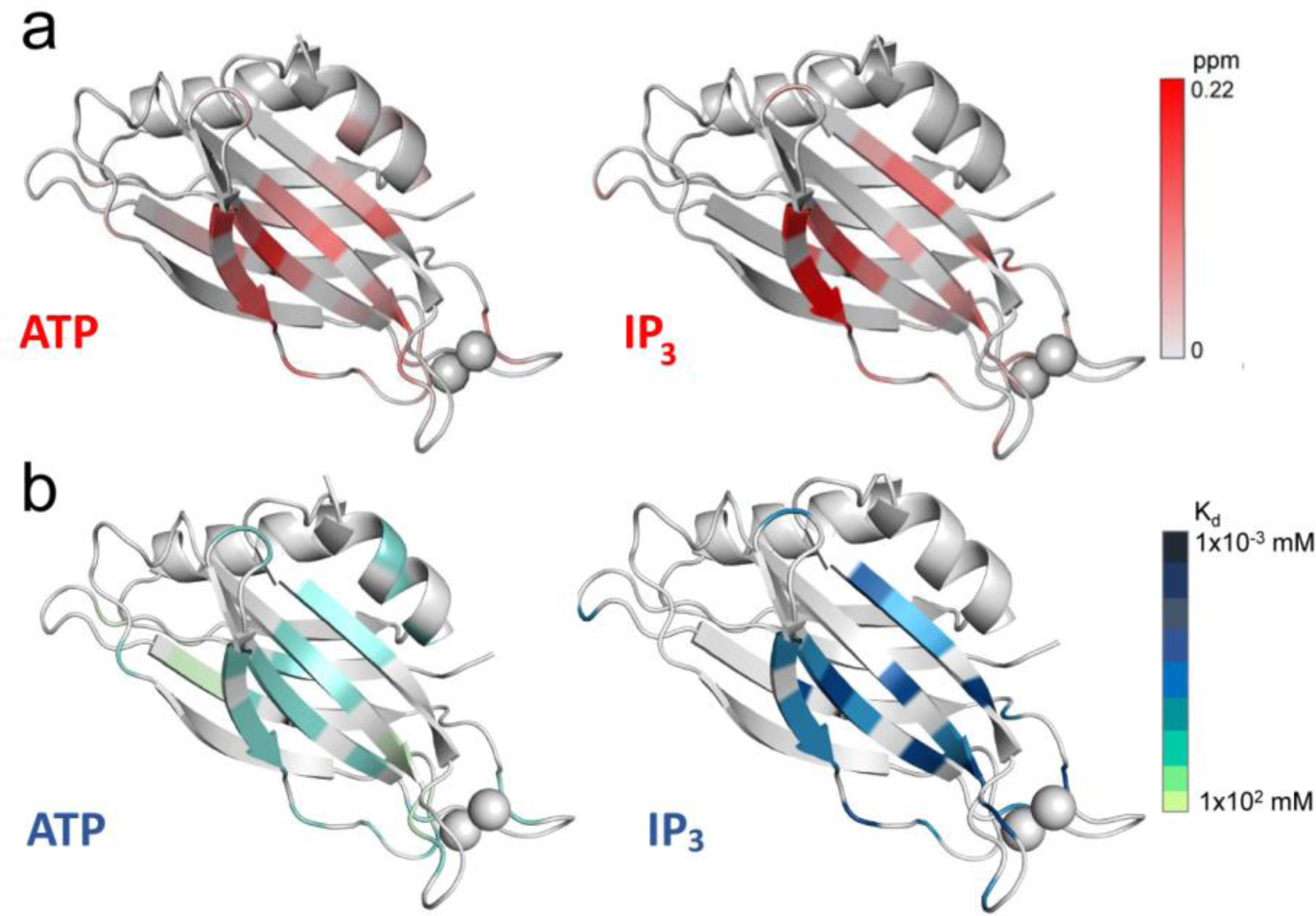
Average weighted chemical shifts (**a**) and affinities (**b**) for ATP and IP_3_ mapped onto the C2B domain of Syt1. Chemical shifts ranged from 0 to 0.22 ppm and binding affinities, K, are given in mM. White ribbon color indicates regions where no contact is detected. IP_3_ and ATP have similar sites of contact in and near the polybasic strand on C2B, although ATP displays a significantly weaker binding affinity than does IP_3_.

### The C2B domain prefers to bind in *trans* to a PIP_2_ containing bilayer, while C2A prefers to bind in *cis* to a PS containing bilayer

The data in Fig. 3 demonstrate that both C2A and C2B have the ability to bridge to membranes containing PS or PIP_2_ when fSyt1 is reconstituted into neutral liposomes. However, a more realistic vesicle lipid composition would include negatively charged lipid in the *cis* membrane, which may inhibit the *trans* binding of the domains. To explore this possibility, we examined the ability of fSyt1 to penetrate membranes when different compositions of charged lipid are used for *cis* versus *trans* vesicles.

The EPR lineshapes from 173R1 in C2A and 304R1 in C2B were shown previously to vary depending upon whether they bound to a PS or PIP_2_ containing bilayer (12). Shown in Fig. 5a are spectra from 173R1 and 304R1 for sC2AB in aqueous solution or when bound to PS or PIP_2_ containing bilayers in the presence of Ca^2+^. The lineshape for 173R1 is significantly different when associated with a PIP_2_ containing membrane than when bound to a PS containing bilayer (see blue and red traces inset Fig. 5a). This difference is due to the fact that site 173 on C2A does not associate as deeply into a PIP_2_ containing bilayer as it does into a PS containing bilayer (a result that was previously verified using power saturation (12)). There is also a slight difference in the lineshape from 304R1 in the 1^st^ Ca^2+^ binding loop of C2B, but this difference is difficult to distinguish in noisier EPR spectra. The lineshape difference for site 173R1 allows us to distinguish whether C2A is associating with a bilayer containing PIP_2_ or PS.

**Figure 5.**
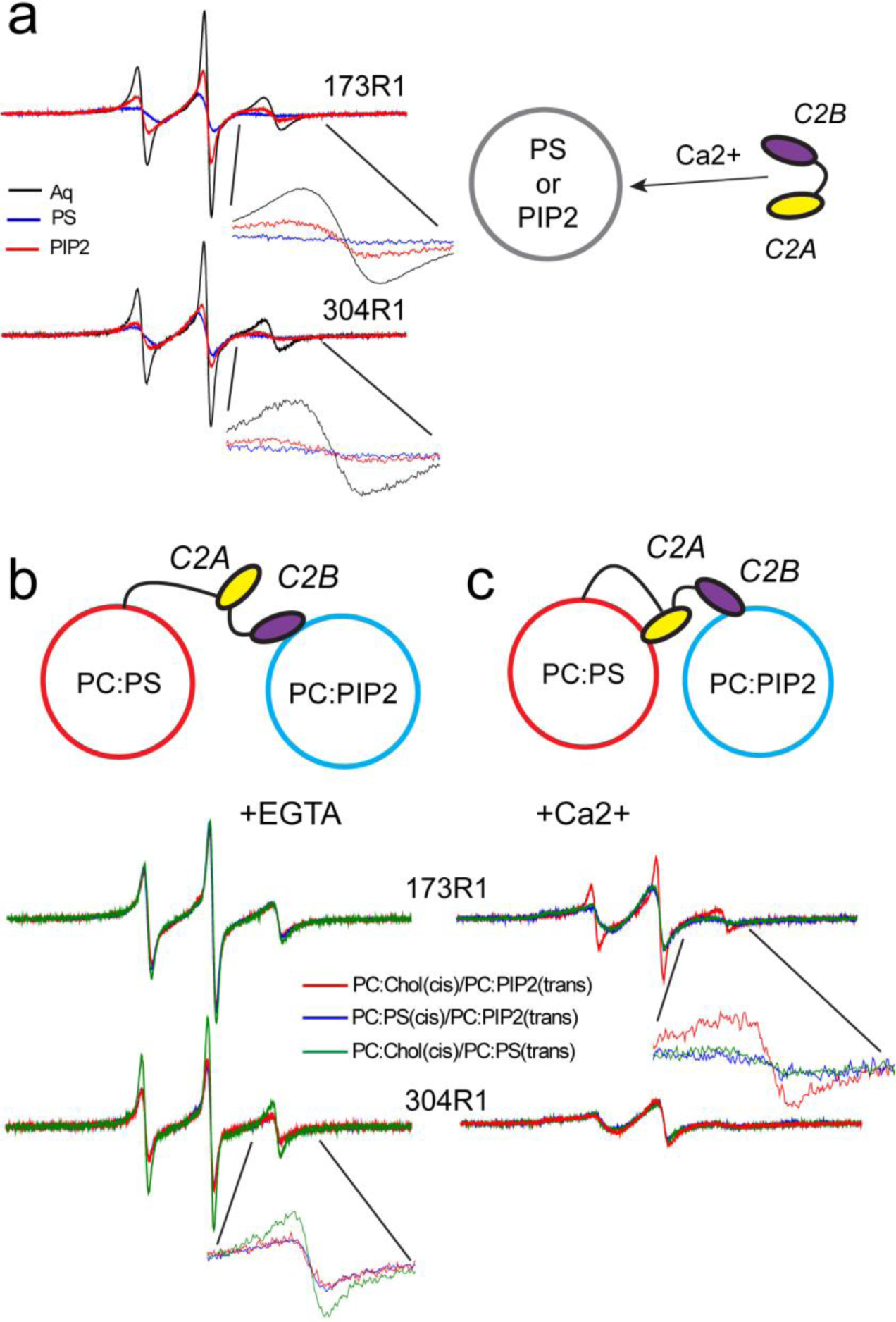
(**a**) EPR spectra from 173R1 or 304R1 in the 1^st^ Ca^2+^-binding loops of the C2A and C2B domains of sC2AB in aqueous solution (black trace), when bound to POPC:POPS (85:15) in the presence of Ca^2+^ (blue trace) or when bound to POPC:PIP_2_ (95:5) bilayers in the presence of Ca^2+^ (red trace). (**b**) EPR spectra from 173R1 and 304R1 in fSyt1 in the absence of Ca^2+^ and (**c**) the presence of Ca^2+^ when fSyt1 is reconstituted into POPC:Chol (80:20) upon the addition of vesicles of POPC:POPS (green trace), POPC:PIP_2_ (red trace); or when reconstituted into POPC:POPS with the addition of vesicles formed from POPC:PIP_2_.

Shown in Figs. 5b,c are EPR spectra obtained from 173R1 and 304R1 when full-length Syt1 is reconstituted into liposomes either with or without negatively charged POPS when vesicles containing either POPS or PIP_2_ are added to the reconstituted fSyt1. As seen in Fig. 5b, in the absence of Ca^2+^ site 173R1 on C2A shows no evidence for membrane insertion when vesicles containing either POPS or PIP_2_ are added to the vesicle mixture. This is consistent with the data in Fig. 3c indicating that there is no membrane insertion of C2A to POPS containing bilayers without Ca^2+^. The spectrum from 304R1 in the 1^st^ binding loop of C2B also shows no evidence for membrane insertion into POPS containing bilayers in the absence of Ca^2+^ (Fig. 5b inset, green trace), however, there is membrane insertion of this site when vesicles containing PIP_2_ are added to reconstituted fSyt1 vesicles (inset, red trace). Calcium-independent insertion is also observed for even when full-length Syt1 is present in a POPC:POPS bilayer (blue trace). The blue and red spectra are identical, and no insertion of C2B into a POPS containing bilayer is observed in the absence of Ca^2+^, indicating that C2B will insert in *trans* into a PIP_2_ containing bilayer, even when POPS is present in the proximal or *cis* membrane. This is consistent with previous measurements showing that the Ca^2+^-independent binding of C2B is much stronger to membranes containing PIP_2_ than membranes containing only PS (12). The plasma membrane is believed to contain high levels of PIP_2_ at the focal site of fusion (51), and these data indicate that C2B will not be free in solution or associated with the vesicle membrane but will preferentially associate with the plasma membrane in the low Ca^2+^ state prior to fusion.

In the presence of Ca^2+^, the C2A domain will associate in *trans* to PIP_2_ containing membranes if Syt1 is present in a neutral membrane. The EPR spectrum for this condition (Fig. 5c, red trace, and also Fig. 3c) resembles that for 173 shown in Fig. 5a in the presence of PIP_2_. It will also associate in *trans* to a POPS containing bilayer if present in a neutral bilayer (green trace). However, C2A will bind in a *cis* configuration if it is reconstituted into a POPS containing membrane even when presented with a PIP_2_ containing vesicle (Fig. 5c, blue trace). Thus, in fSyt1 C2A prefers a POPS containing bilayer over a PIP_2_ containing bilayer. For C2B, the spectra in Fig. 5c show that site 304R1 on the C2B domain undergoes *trans* interfacial insertion into either a POPS or PIP_2_ containing bilayer in the presence of Ca^2+^ when reconstituted into POPC:Chol. Because the EPR lineshapes from site 304R1 when bound to PIP_2_ or POPS are very similar (inset, Fig. 5a), it is not possible to distinguish whether C2B is binding in *cis* or *trans* when fSyt1 is reconstituted into a POPS containing membrane. The fact that C2B is attached in *trans* to the PIP_2_ containing bilayer in the absence of Ca^2+^ and the dramatically higher affinity of C2B for the PIP_2_ containing bilayer suggests that that C2B will remain attached to and insert into the PIP_2_ containing bilayer with increases in Ca^2+^. Since PIP_2_ is believed to reside primarily in the plasma membrane, the two domains are likely to bind to opposing bilayers.

### Pulse EPR data indicate that well-defined orientations between C2A and C2B are not present in full-length Syt1

Distance measurements using double electron-electron resonance (DEER) were previously used to examine the relative orientations of C2A and C2B in the sC2AB fragment and demonstrated that a bridging of sC2AB between two bilayers by the domains was a likely orientation for the fragment (23). Here we made several measurements to examine the relative orientations between C2A and C2B in the full-length protein under conditions that the data in Fig. 5 predict should encourage either a *cis* or *trans* binding of the C2A and C2B domains.

Shown in Fig. 6a are DEER data obtained between sites 173R1 and 323R1 in C2A and C2B, respectively. As seen previously in sC2AB, the distributions obtained for the fSyt1 are broad, and were fit in the present case with a single Gaussian. Other data obtained between labels in C2A and C2B in the full-length protein also yield broad distances (Fig. S4). As found previously for the sC2AB domain, the data indicate that the C2 domains of Syt1 are flexibly linked and do not assume a well-defined orientation relative to each other in the full-length protein (23).

**Figure 6.**
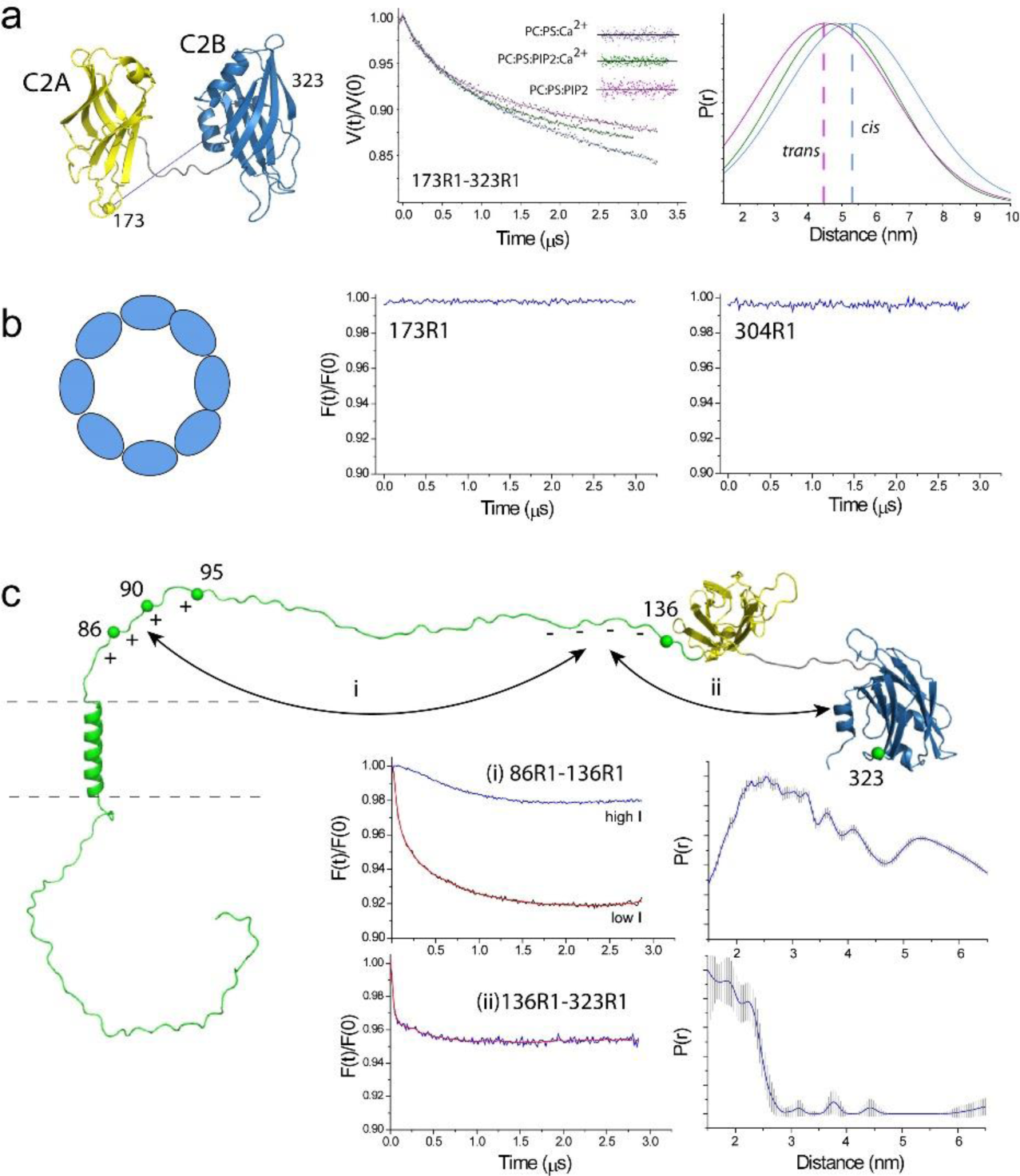
Pulse EPR on fSyt1. (**a**) Raw DEER data obtained for the 173R1/323R1 spin pair under conditions where fSyt1 is reconstituted into POPC:POPS with bound Ca^2+^ (blue trace), with bound Ca^2+^ and the addition of PIP_2_ containing vesicle (green trace) and without Ca^2+^ and the addition of PIP_2_ containing vesicles (magenta trace). The data were well fit with single Gaussian distributions. (**b**) Background corrected DEER data obtained for the single labels 173R1 and 304R1 for fSyt1 reconstituted into POPC:Chol in the absence of Ca^2+^. There is no indication of a dipolar interaction between labels in these samples, which indicates aggregation of Syt1 is not occurring within this membrane environment. (**c**) DEER data for the 86R1-136R1 spin pair (interaction *i*, see text) of fSyt1 reconstituted into POPC:Chol taken under low (red trace) and high (blue trace) ionic strength conditions. Also shown are background corrected DEER data for the 136R1-323R1 spin pair (interaction *ii*), which indicate the presence of a transient interaction between the polybasic face of C2B and the negatively charged segment in the linker.

The distances between C2A and C2B in the full-length protein were explored under conditions that should favor either a *cis* binding of both domains (blue trace), under conditions where only the C2B domain should bind in *trans* to a PIP_2_ vesicle (magenta trace) and where a bridging or a mixture of *cis* and *trans* states should be present (green trace). This is expected to change the relative orientations of the domains and hence the interspin distances between these labeled sites. As seen in Fig. 6a, the shortest average distance was obtained under conditions where C2B is attached in *trans* or when there is likely to be a bridging between two vesicles. The longest distance was obtained when C2A and C2B should both be bound in a *cis* configuration (blue). This is consistent with the distances expected between sites 173 and 323 when the domains are oriented either parallel or antiparallel to each other. Moreover, the distance difference between these two configurations is expected to be about 9 Angstroms, which approximately the difference seen in Fig. 6a.

### Oligomerization of the C2 domains of full-length Syt1 cannot be detected

It should be noted that for fSyt1, no dipolar interactions can be detected for single labeled sites in C2A or C2B, and the distance distributions shown in Fig. 6a are not a result of oligomerization. Figure 6b shows the background corrected DEER data obtained for 304R1 and 173R1 in the full length protein in the absence of Ca^2+^ when reconstituted into POPC:Chol at a relatively high protein:lipid ratio of 1:200. No significant dipolar interactions are detected for this single labeled site. Dimers or ring-like oligomers of Syt1 should produce signals corresponding to inter-spin distances of approximately 40 Angstroms, which is well-within the range that can be distinguished here. The absence of any signal in this case indicates that oligomers are not significantly populated. If the protein were simply dimerized, the signal-to-noise in these data indicate that less than 5% of the protein would be present in this state. A number of conditions were examined, and no evidence for oligomers could be detected by EPR using the full-length protein (see Fig. S5).

### Intramoleular interactions in the linker are weak and driven by electrostatics

It has been proposed that oppositely charged segments on the juxta membrane linker might interact electrostatically, thereby regulating the effective length of the linker (52). This interaction, labeled (i) in Fig. 6c, was examined in fSyt1 by placing labels within these two regions at sites 86 and 136. The modulation depth of the DEER signal obtained for 86R1/136R1 at low ionic strength (red trace) is 8%, which is less than half of that expected for a stable interaction between these regions under the conditions of this experiment. As seen in the blue trace, the signal was virtually eliminated by running the experiment at higher ionic strength. Although there is clearly an interaction between these domains in the full-length protein, these data indicate that the interaction is transient and electrostatic in origin.

Another interaction that can be detected is an interaction between the highly positively changed face of C2B with the negatively charged region in the linker, labeled (ii) in Fig. 6c. The modulation depth resulting from this dipolar interaction is 4%, and is also much less than that expected for a stable intramolecular interaction. This suggests that this interaction, like the interaction between oppositely charged segments of the linker, is weak and transient. We also ran control experiments under these conditions to insure that single labeled sites in the linker did not give rise to strong DEER signals due to oligomerization. These data, as well as EPR spectra from the linker, suggest that the linker is largely monomeric, unstructured, and that only small portion of the linker adjacent to the transmembrane segment is associated with the membrane interface (see Fig. S6 and Table S2).

## Discussion

Synchronous neurotransmitter release is mediated by Ca^2+^-binding to the C2A and C2B domains of synaptotagmin 1. It is not presently clear how this binding event triggers the assembly of the SNAREs and the fusion of the synaptic vesicle with the plasma membrane. The Ca^2+^-dependent membrane insertion of the C2A and C2B domains is robust and persists under conditions where the Syt1-SNARE interactions are weak or absent. For example, normal ionic strength and normal cellular levels of ATP/Mg^2+^ largely eliminate Syt1/SNARE interactions, which appear to be weak and heterogeneous. However, under the same conditions, the binding of Syt1 to bilayers containing PIP_2_ persists (19). As a result, a number of models have focused on the membrane interactions of the C2 domains Syt1 as the mediators of membrane fusion.

The polybasic face of C2B is an important structural feature of Syt1, and it determines much of the behavior of the protein. It has been known for some time that isolation of C2B or sC2AB requires an additional ion exchange step to remove acidic contaminants from the protein, and that these contaminants will eliminate the membrane binding of C2B and promote aggregation of the domain (50). Charge mutations in the polybasic face of C2B dramatically alter the affinity of the domain towards membranes containing PIP_2_ (12) and also have effects on fusion *in vivo* (14). Here, the presence of polyelectrolytes such as ATP at normal cellular levels is found to interfere with the membrane binding of sC2AB (as shown in Fig. 2) and the membrane insertion of fSyt1 (as shown in Fig. 3), however the presence of PIP_2_ in the membrane can overcome this inhibition towards binding and insertion. This is consistent with the data presented in Fig. 4, which show that the PIP_2_ headgroup has a much higher affinity for the polybasic face of C2B than does ATP. These results are also consistent with previous work using two-color confocal microscopy showing that vesicles containing Syt1 will not attach to opposing vesicles in the presence of ATP unless PIP_2_ is present in the target membrane (44).

The data presented here examined the *cis* versus *trans* interactions of the C2A and C2B domains in membrane-reconstituted full-length Syt1. The ability to bind in *trans* appears to be due in part to the polybasic face of C2B and the lipid preferences of the two domains. In the absence of Ca^2+^, the C2A domain does not penetrate charged membrane interfaces, but the C2B domain will penetrate to an opposing lipid bilayer containing PIP_2_ even when Syt1 is reconstituted into a membrane containing POPS. This finding differs somewhat from results obtained using a confocal vesicle tethering assay where no bridging is detected in the absence of Ca^2+^ (53). This difference is likely due to the fact that the present experiments are carried out at much higher accessible lipid concentrations, which would favor membrane association for the weaker Ca^2+^-independent binding mode of C2B. At the focal site of fusion, the plasma membrane is believed to achieve high levels of PIP_2_ during the priming events for fusion, perhaps in excess of 5% (51); and the local lipid concentrations are likely to be high. As a result, the C2B domain is expected to be bound to the plasma membrane in the pre-fusion state. In the presence of Ca^2+^, the C2A domain binds preferentially to PS containing bilayers and it penetrates preferentially into a *cis* membrane containing PS even when presented with lipid vesicle containing PIP_2_.

The data presented here indicate that the long linker connecting C2A to the transmembrane helix in the synaptic vesicle is largely monomeric and unstructured. In previous work, the long linker connecting C2A and C2B to the membrane interface was reported to oligomerize (54). However, oligomerization of full-length Syt1 is dependent upon the detergent used for protein reconstitution. In the protocol employed here, the protein was reconstituted from CHAPS, which produces EPR spectra from the C2A and C2B domains that resemble those obtained for the sC2AB fragment (29). Using this protocol, minor levels of oligomerization within the linker region are observed. There is also no significant dimerization or oligomerization of the C2A or C2B domains of fSyt1 when membrane reconstituted. This suggests that the tendency of Syt1 to oligomerize in the full-length protein is at best very weak, a finding that is not consistent with models where rings of oligomerized Syt1 provide a steric block to vesicle-plasma membrane fusion in the absence of Ca^2+^ (26).

When reconstituted using CHAPS, the only region of the linker that appears to be in membrane contact is the segment close to the transmembrane vesicle anchor (Table S2). Acylation may be present within the linker (55); however, the cysteines that may be modified lie in this membrane associated region are close to the transmembrane anchor, and they would not be expected to alter the C-terminal side of the linker. As a result, the majority of the linker encompassing perhaps as many as 50 residues is likely to lie in the aqueous phase in an unstructured state. Although we do not know the state of the SNAREs or their associated regulatory proteins prior to fusion, the long linker would allow the C2A and C2B domains to diffuse a significant distance from the vesicle membrane interface in the pre-fusion state. As shown in Fig. 7, this may allow C2B to interact with the PIP_2_ containing plasma membrane while C2A remains free in the cytosol. This arrangement gives Syt1 the ability to regulate the vesicle-plasma membrane distance upon the addition of Ca^2+^. The addition of Ca^2+^ would insert the C2 domains and limit the plasma membrane-vesicle membrane distance to the dimension of the C2 domains (about 4 to 5 nm). This is in fact consistent with a fluorescence based lifetime measurement indicating that Ca^2+^ reduces membrane separation by 2 to 3 nm to a distance of about 5 nm (25).

**Figure 7.**
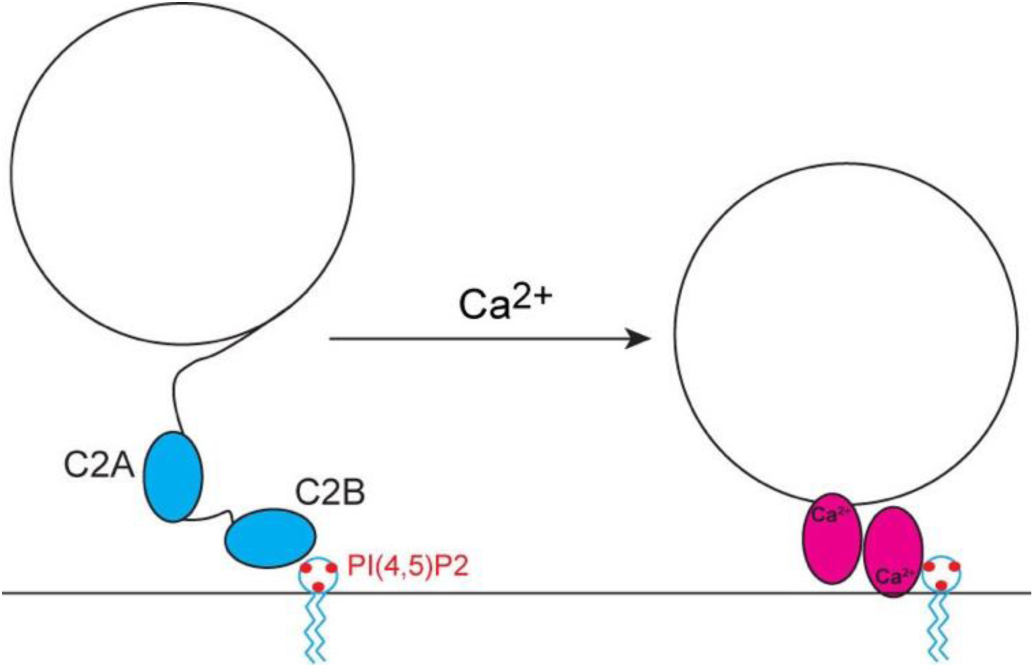
The lipid affinities of C2A and C2B and the long linker connecting C2A to the synaptic vesicle suggest that Syt1 may be configured to regulate the vesicle-plasma membrane distance upon the addition of calcium.

The back-to-back binding of the C2A and C2B domains to opposing bilayers such as that shown in Fig. 7 was proposed previously based upon EPR restraints (23) and later based upon NMR data (24). It also appears to be a feature of the C2B domain, which may bridge bilayers on its own due to a highly positively charged region that lies opposite the Ca^2+^-binding site (56). There is also support for this type of model in earlier work proposing that PIP_2_ directs Syt1 towards the plasma membrane (57) and appears to be required in the target membrane for fusion activity (58, 59).

Exactly how changing the intermembrane vesicle-plasma membrane distance might trigger fusion is unclear. The pre-fusion complex is likely to involve a number of proteins, including the t-SNAREs in a prefusion state with Munc18 (60), complexin (61) and perhaps synaptobrevin. Synaptobrevin might either be partially associated to the N-terminal region of the t-SNAREs or be complexed with Munc18, as has been suggested for yeast SNAREs (62). This is likely to be a very crowded environment, and the shortening of this inter-membrane distance may bring the appropriate components into contact to trigger SNARE assembly, or might remove steric constraints on the assembly of the complex.

## Author contributions

S.B.N. and D.S.C. wrote the paper. S.B.N, A.T. and D.S.C. designed the experiments. S.B.N and A.T. carried out the work.

## Acknowledgements

We would like to acknowledge the support of NIH grant PO1 GM072694 (to D.S.C.). We would like to thank Profs. Reinhard Jahn, Lukas Tamm and members of their research groups for helpful discussions during the course of this work. We would also like to acknowledge David Nyenhuis for help acquiring and processing DEER data and Vanessa Bajik for assistance with the NMR experiments. The authors declare that they have no financial conflict of interest with this work.

